# Establishing a cell-free *Vibrio natriegens* expression system

**DOI:** 10.1101/331645

**Authors:** Daniel J. Wiegand, Henry H. Lee, Nili Ostrov, George M. Church

## Abstract

The fast growing bacterium *Vibrio natriegens* is an emerging microbial host for biotechnology. Harnessing its productive cellular components may offer a compelling platform for rapid protein production and prototyping of metabolic pathways or genetic circuits. Here, we report the development of a *V. natriegens* cell-free expression system. We devised a simplified crude extract preparation protocol and achieved >260 μg/mL of super-folder GFP in a small-scale batch reaction after three hours. Culturing conditions, including growth media and cell density, significantly affect translation kinetics and protein yield of extracts. We observed maximal protein yield at incubation temperatures of 26°C or 30°C, and show improved yield by tuning ions crucial for ribosomal stability. This work establishes an initial *V. natriegens* cell-free expression system, enables probing of *V. natriegens* biology, and will serve as a platform to accelerate metabolic engineering and synthetic biology applications.

---

With the shortest doubling time of all known organisms, *Vibrio natriegens* has garnered considerable interest as a promising microbial host to accelerate research and biotechnology ^1–4^. Its rapid growth rate, which has been linked to high rates of protein synthesis ^5^ and metabolic efficiency ^6,7^ suggests that this host may be harnessed as a powerful cell-free expression system. Cell-free bioproduction of protein and chemicals has been extensively investigated in *E. coli* and recently expanded to several other bacteria ^8–13^. Accordingly, we assessed *V. natriegens* crude cell extract productivity in small-scale batch reactions using super-folder GFP (sfGFP) controlled by a T7 promoter. We systematically explored culturing conditions and extract additives to provide a simple and robust *V. natriegens* cell-free system. Our calibration offers initial conditions for cell-free protein expression with *V. natriegens* and sheds light on critical factors for future process engineering ^14,15^

To begin developing a cell-free protocol for *V. natriegens*, we evaluated established methods for extract preparation for *E.coli* and other bacterial systems ^10,16–18^. Overall, we aimed for general accessibility and reproducibility by opting at each step for commonly available equipment while considering ease of user operation and cost reduction. Our preparation protocol utilizes one liter cultures in shake flasks, cell lysis by pulse sonication, and small-scale (10 μL) batch reactions in a 96-or 384-well format to maximize parallelization and screening throughput ^17^.

We first assessed the effect of *V. natriegens* culturing conditions on extract productivity. We tested commonly available microbiological media, supplemented with salt as needed to maintain robust growth ^1,2,5,19^. Crude cell extracts were prepared from *V. natriegens* cultures growing in the following six media: LB with 3% (w/v) NaCl (LB3), LB with V2 salts (LB-V2), LB with Ocean Salts (LBO), Nutrient Broth with Ocean Salts (NBO), Brain Heart Infusion Broth with Ocean Salts (BHIO), and Marine Broth (MB). Cultures were grown at 30°C and harvested at OD_600_= 1.0 (**Methods**).

We found that the choice of growth media significantly affects cell extract productivity. Extracts cultured in LB-V2 produced the highest protein yield (196 ± 12.46 μg/mL), followed by extracts cultured in BHIO (58.86 ± 2.61 μg/mL) and LB3 (33.18 ± 5.57 μg/mL) (**Figure 1a, Supplementary Figure 1a**). Low protein yield was observed from extracts cultured in NBO and LBO media, with no significant protein expression observed from extracts cultured in MB media. We thus elected to use LB-V2 extracts for all further investigations.

**Figure 1.**
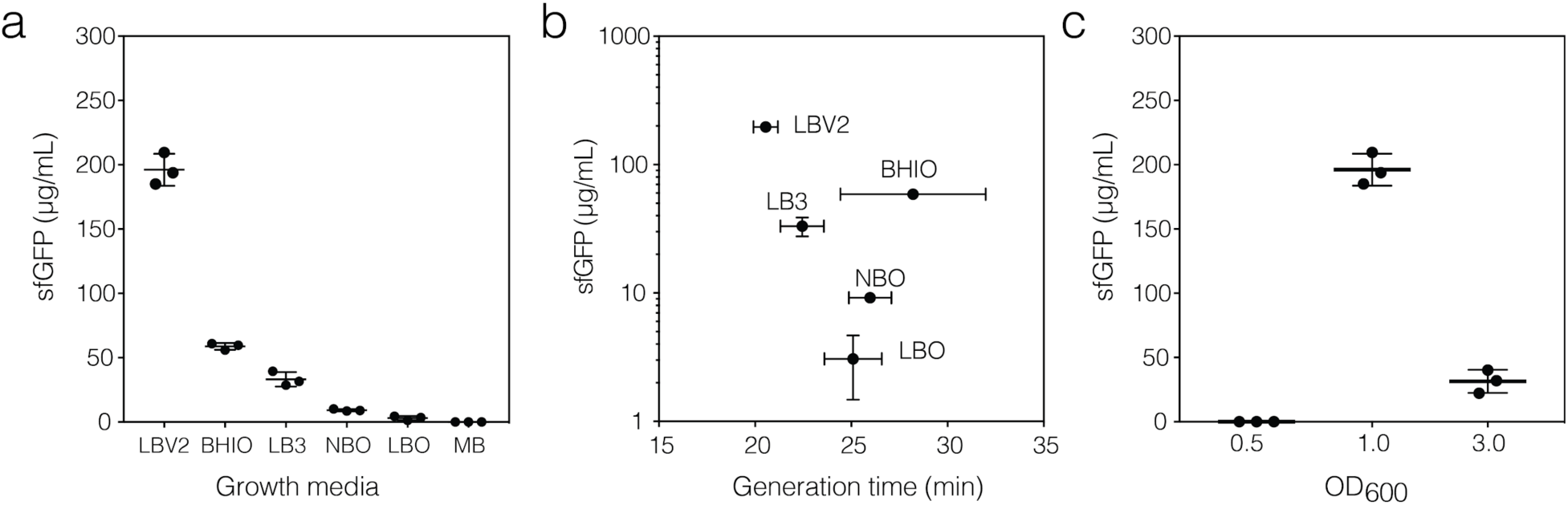
*V. natriegens* culture conditions impact extract protein yield. (a) Effect of culture growth media on cell-free extract protein yield. *V. natriegens* cells were grown in the indicated media at 30°C to OD_600_ = 1.0. Crude cell lysate was prepared production of sfGFP measured over three hours. (LB3 - LB with 3% (w/v) NaCl; LB-V2 - LB with V2 salts; LBO - LB with Ocean Salts, NBO - Nutrient Broth with Ocean Salts; BHIO - Brain Heart Infusion with Ocean Salts; and MB - Marine Broth). (b) Generation time and sfGFP yield for each media type tested. (c) Optical density at the time of cell harvest. Unless otherwise indicated, cell-free reactions were incubated at 26°C in a thermocycler using 500 ng of plasmid DNA, 160 mM K-glutamate and 3.5 mM Mg-glutamate. sfGFP yield was measured after 3 hours. The mean and standard deviations are shown (N=3).

Previous studies have suggested that higher cellular ribosomal content is required to support faster growth rates ^5,20^. To test whether extract productivity could be used for a proxy for active ribosomes in each growth medium, we assessed the correlation between the culture generation time and extract protein yield (**Figure 1b, Supplementary Figure 1c**). Our data shows no relationship between generation time and cell extract productivity. The three shortest generation times at at 30°C were MB, LB-V2, and LB3 (20.14 ± 2.16, 20.55 ± 0.63 and 22.14 ± 1.13 minutes, respectively). However, extracts from LB-V2 cultures produced significantly more protein than extract from all other culture media (∼6-fold more than LB3).

To gain deeper insight into the difference between growth media, we assessed translation kinetics for each growth media type by examining the rate of accumulating sfGFP (**Figure S1b, Supplementary Figure 1b, Methods**). As expected, LB-V2 extracts had the maximum rate of protein synthesis, which was achieved after ∼30 minutes. However, NBO extracts reached its maximum synthesis rate in only ∼15 minutes, although the rate was 2.6-fold lower, and the total protein yield was 19.6-fold lower. Intriguingly, although their rate and total yield were significantly lower than that of LB-V2 extract, BHIO and LB3 extracts appear to sustain protein synthesis more robustly compared to other extracts, as evidenced by the protracted time constants for maximum and minimal rates. All extracts demonstrated a decay in rate of protein expression after 60 minutes, indicating consumption of input building blocks, buildup of inhibitory byproducts, or depletion of the sfGFP template. Overall, these results indicate significant variation in the fraction of active translation between media types. Specifically, potassium and magnesium ions, which are present in LB-V2 but not LB3, may enhance the stability of translation components in crude cell extract. Given the significant impact of culture media on extract yield, further investigation and development of customized cell-free growth medium for extract preparation may be required.

Additionally, we tested whether a post-lysis run-off reaction, previously shown to increase the availability of translational machinery in crude cell extracts ^17^, would be beneficial to increasing protein yield. We performed run-off reactions prior to addition of the sfGFP template, as previously reported ^21,22^. We found that run-off reactions resulted in a significantly reduced capacity to produce sfGFP (**Supplementary Figure 8**).

Active cell-free extracts are routinely produced using exponentially growing cells ^17,23^. We thus tested the impact of *V. natriegens* cell density at harvest on extract productivity. Cells were grown in LB-V2 media, harvested at OD_600_ of 0.5, 1.0 or 3.0 and the corresponding sfGFP yield was measured (**Figure 1c, Supplementary Figure 2**). We found the most productive extracts were obtained from mid-logarithmic cultures (OD_600_ = 1.0), compared to 6.2-fold lower yield from late-logarithmic cultures (OD_600_ = 3.0). We were unable to detect sfGFP production from extracts obtained from early-logarithmic cultures (OD_600_ = 0.5). These effects of cell density on extract performance are consistent with those reported for other cell-free systems ^17^.

Cell-free extracts uniquely allow cellular components to operate decoupled from their culturing conditions. We next turned to calibrating cell-free reaction conditions. We examined incubation temperatures ranging 18-37°C and observed maximal protein expression at 26°C and 30°C (262.28 ± 8.13 μg/mL and 265.01 ± 8.15 μg/mL, respectively) (**Figure 2a**). While *in vivo* growth rate for *V. natriegens* has been found to increase with temperature and to be maximal at 37°C ^1,19^, we found that extracts incubated at 30°C produced 3-fold more protein than 37°C. These results support the hypothesis that the *V. natriegens’* rapid growth at a higher temperature is aided by a larger number of ribosomes, rather than more efficient ones ^5^. It may also be the case that more functional ribosomes are present at lower temperatures despite a higher total number at higher temperature as previously observed in *E. coli* ^24^. Further investigation of ribosome content and availability is warranted to elucidate underlying biology. In addition, the reduced protein yields at temperatures higher than 30°C may be artifactual due to our choice of T7 polymerase. Due to its high processivity, T7 polymerase has been shown to reduce protein yield for a bacterial cell-free system by disrupting the coupling of transcription and translation ^25^. Further improvements may be made by using mutant T7 polymerases or endogenous promoters. For ease of further investigations, we chose to incubate all *V. natriegens* cell-free reactions at 26°C.

**Figure 2.**
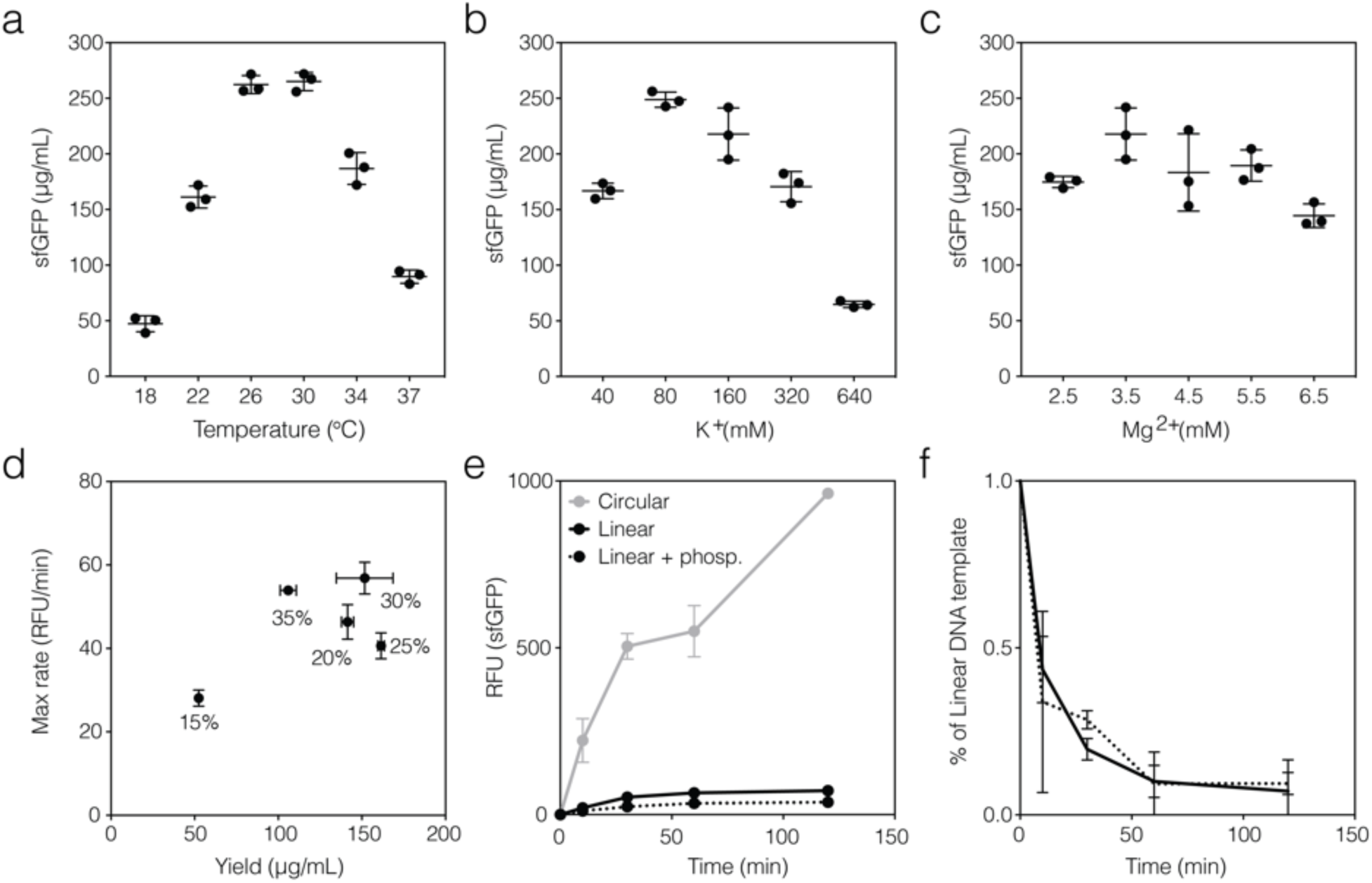
Properties of *V. natriegens* crude cell-free extract. (a) Cell-free incubation temperature. (b) Supplemented potassium ions in reaction buffer. (c) Supplemented magnesium ions in reaction buffer. (d) Percent of extract used relative to total reaction volume. (e) Template DNA provided in the cell-free reaction. Equimolar amount of circular plasmid (pJL1, grey), linear DNA (PCR product, black solid line) or linear DNA with two phosphorothioated bonds on each end (PCR product, black dotted line) was used as template for cell-free expression of sfGFP. (f) Degradation of linear DNA template. Fluorescence of AlexaFluor 594-5-dUTP labeled DNA template was monitored over two hours. Linear template with (dotted) or without (solid) two phosphorothioated bonds on each end was used. Unless otherwise indicated, all experiments were performed using *V. natriegens* crude cell extract incubated at 26°C for three hours, supplemented with 80 mM K-glutamate and 3.5 mM Mg-glutamate. The mean and standard deviations are shown (N=3).

Having set an incubation temperature for optimal extract productivity, we then tuned the range of K^+^ and Mg^2+^ ions in the extract reaction ^10,18^. In general, we found the addition of K^+^ had a significant effect on both yield and rate of protein expression, while Mg^2+^ had only moderate effect at the tested conditions (**Figure 2b-c, Supplementary Figure 3**). We set our assay conditions at 80 mM K^+^ and 3.5 mM Mg^2+^, which yielded the highest reaction productivity for sfGFP expression (248.75 ± 6.82 μg/mL and 217.87 ± 23.40 μg/mL, respectively). Notably, while higher ion concentrations have been reported ^26^, the determined optimal ion concentrations for both K^+^ and Mg^2+^ in our system are consistent with other cell-free systems which have been developed with 3-Phosphoglyceric acid (3-PGA) as an energy source ^10,18,27^.

We then sought to determine if the total amount of crude extract needed per sample could be lowered while retaining reaction productivity. The percent of cell-free reaction that comprises crude cell extract varies between different bacterial systems ^12,17^ and titration of the cellular components has been shown to improve protein yield ^9^. We thus tested reactions where the percent of extract was varying, ranging 35% to 15%, and monitored the kinetics of sfGFP accumulation. Surprisingly, we found that decreasing the extract from 35% to 30% of the total reaction volume resulted in significant improvement in protein yield (43% increase) with no significant change in expression rate (**Figure 2d**, **Supplementary Figure 4**). Further decrease in extract amount (from 30% to 25%) resulted in a decreased rate of protein expression without significant change in protein yield. These results indicate up to half of the original extract volume can be used with increased or equal reaction productivity, which increases the number of cell-free reactions produced from a single culture batch. It is worth also noting that the lowering the total extract percentage per reaction volume yielded sustained protein production for the duration of the reaction (**Supplementary Figure 4**).

The use of linear, rather than plasmid, template DNA in cell-free expression systems is desirable for high-throughput and rapid testing ^18^. We thus evaluated protein production from a linear PCR template compared to a circular plasmid template (**Figure 2e**). In addition to sfGFP under T7 promoter, our PCR amplicon also contained 58bp of non-coding DNA at the 3’ end of the construct. We observed protein production from the PCR amplicon to be 13.5-fold lower than an equimolar amount of circular plasmid, suggesting rapid DNA degradation by cellular nucleases. To test whether a linear template could be protected from nuclease digestion, we assessed protein yield from a PCR product with two phosphorothioated bonds on each end ^24^. Notably, this modified template was also 58bp shorter, carrying less non-coding sequence at the 3’. We observed no improvement in protein yield using protected PCR amplicon (**Figure 2e**), suggestive of endonuclease, rather than exonuclease, activity. The modified short template had 2-fold lower yield compared with longer, non-modified template. Padding of linear templates with non-coding sequences ^28^ or inactivation of endogenous nucleases by genome engineering will likely improve protein yield significantly and warrants further investigation.

To monitor the kinetics of linear DNA template degradation, we used a fluorescence-based assay to quantify the amount of intact linear DNA template in the cell-free reaction over time (**Methods**) ^28^. We found that more than 50% of linear DNA template was degraded by 10 minutes of co-incubation with the cell-free extract (**Figure 2f**). No significant difference in degradation was observed using the shorter phosphorothioated template, corroborating our previous observations that protecting the 5’ ends of linear DNA does not improve template stability and that less non-coding padding decreases protein yield.

Previously, it was shown that linear template stability can be increased in *E.coli* cell-free system with inclusion of the lambda phage protein GamS, which inhibits the host RecBCD nuclease complex ^28,29^. However, we observed no change in the extent of linear template degradation upon addition of lambda GamS protein to *V. natriegens* cell-free reactions (**Supplementary Figure 9)**. The same results were observed for both phosphorothioate protected and unprotected linear templates. This could indicate that GamS does not interact with the DNA-binding domain of *V. natriegen*s’ RecBCD exonuclease complex or that linear template degradation occurs by an independent mechanism. Alternatively, degradation may result from endonuclease activity given that the inclusion phosphorothioate bonds at both the 5’ and 3’ termini the linear template did not alter degradation kinetics (**Figure 2f**). Additional research is required to elucidate *V. natriegens* exo-or endonuclease and effective strategies to inhibit their activity. Engineering of the host genome to remove nucleases may enable facile production of highly active extracts for high efficiency protein expression with linear templates. ^30^.

Maximizing protein expression in cell-free systems requires the combinatorial optimization of a large parameter search space and no single best approach currently exists ^21,22,31,32^. During this work, an alternative *V. natriegens* CFPS has been reported ^26^, which explores the use of other culture medium, energy regeneration sources, and extract preparation. For example, while our screen of growth media shows that the use of LB-V2, rather than BHIO, results in higher protein expression yields (**Supplementary Figure 1**), the alternative report showed that an extensive purification process can provide more cell-free protein expression in cells harvested from BHIO culture media at high optical density (OD=2.5). Additionally, we found that run-off did not promote increased protein expression, which may indicate altered ribosomal occupancy under different culture conditions or growth stage at the time of harvest. Consistent with previous reports ^33–35^, it is interesting to note the higher concentration of Mg^2+^ required when using creatine phosphate (CP) as energy regeneration system compared with 3-PGA in this work. Additional investigation will be essential to fully develop the potential of *V. natriegens* for CFPS. While protein yield is an important feature, it may not be the sole consideration when working with CFPS. The protocol for extract preparation presented in this work does not require specialized equipment or lengthy purification steps, and is therefore short, accessible, and simple to execute, which should facilitate further experimentation.

Throughout the development of this preparation protocol for *V. natriegens* CFPS, we have utilized the same methods and reagents, where appropriate, to prepare extracts from *E. coli* A19, an RNase I-deficient strain that is commonly used for cell-free reactions, to validate protein expression functionality ^36,37^. Crude cell extract was prepared as previously described from *E. coli* cultured in rich media (YPG) at 37°C ^23^ and harvested at OD_600_=1.0 (**Methods**). We then performed similar calibration of reaction conditions for *E.coli* as described above for *V. natriegens* (**Supplementary Figure 5**). For context, we observed comparable protein yields between *E.coli* A19 and wild-type *V. natriegens* (**Supplementary Figure 6**) when production occured in their respective optimal reaction conditions. The yield obtained from *E. coli* A19 extracts are similar to previously reported GFP yields using small-batch conditions ^38^, which supports the robustness of our protocol. Kinetic analyses indicated that *V. natriegens* extracts sustained elevated protein expression rates over 60 minutes, while *E. coli* extracts only sustain elevated rates for 20 minutes. More broadly, this system produces more protein than cell-free systems derived from *Bacillus subtilis* ^27^ and *Streptomyces venezuelae* ^10^, as measured by sfGFP yield (∼9.3 μM sfGFP in our 10 μL *V. natriegens* cell-free reaction, compared with 0.8 μM and 1.3 μM, respectively).

Taken together, these results provide a foundation for cell-free protein expression with wild-type *V. natriegens* and may direct further efforts to establish designated cell-free strains. Higher protein yield could be achieved by further optimization of culturing and preparation conditions and may include: semi-continuous extract reactions to allow for energy regeneration, resupplying of amino acids, and the removal of waste products ^12,39^. In addition, engineering of *Vibrio natriegens,* for example using DNAse-or RNAse-deficient strains, removal of deleterious and competing metabolic pathways, and expression of additional tRNAs will likely further enhance protein yield from extracts ^40^. These additional improvements will also enable the use of linear DNA to facilitate rapid prototyping and high-throughput screens. This work establishes an initial cell-free system in *V. natriegens* and sheds light on important aspects for continued development.

## Supporting Information

Attached supporting information document includes all materials & methods and supplementary figures for the main text.

## Author contribution

DJW, NO, and HHL designed and performed experiments. DJW, NO, and HHL analyzed data and wrote the manuscript. GMC supervised the study.

## Acknowledgments

The authors would like to thank Nina Donghia for helpful discussion regarding cell-free extract preparation and Dr. David Thompson for providing sfGFP standards.

## Notes

DJW, NO, and GMC have filed a patent related to this work.

